# *ALTERED MERISTEM PROGRAM1* regulates leaf identity independent of miR156-mediated translational repression

**DOI:** 10.1101/856864

**Authors:** Jim P. Fouracre, Victoria J. Chen, R. Scott Poethig

## Abstract

In *Arabidopsis*, loss of the carboxypeptidase, ALTERED MERISTEM PROGRAM1 (AMP1), produces an increase in the rate of leaf initiation, an enlarged shoot apical meristem and an increase in the number of juvenile leaves. This phenotype is also observed in plants with reduced levels of miR156-targeted *SQUAMOSA PROMOTER BINDING PROTEIN-LIKE* (*SPL*) transcription factors, suggesting that AMP1 may promote SPL activity. However, we found that the *amp1* phenotype is only partially corrected by elevated *SPL* gene expression, and that *amp1* has no significant effect on *SPL* transcript levels, or on the level or the activity of miR156. Although evidence from a previous study suggests that AMP1 promotes miRNA-mediated translational repression, *amp1* did not prevent the translational repression of the miR156 target, *SPL9*, or the miR159 target, *MYB33.* These results suggest that *AMP1* regulates vegetative phase change downstream of, or in parallel to, the miR156/*SPL* pathway and that it is not universally required for miRNA-mediated translational repression.

**Summary statement:** We show that loss of the carboxypeptidase, AMP1, does not interfere with the function of miR156 or miR159, suggesting that AMP1 is not universally required for miRNA-mediated translational repression in *Arabidopsis*.

## INTRODUCTION

Plant life-histories are underpinned by a series of developmental transitions, the correct timing of which are crucial to plant survival and reproductive success (Huijser and Schmid, 2011). Vegetative phase change describes the switch between the juvenile and adult stages of vegetative growth. Depending on the species, this transition can lead to shifts in a wide variety traits (Poethig, 2013). In the model plant, *Arabidopsis thaliana*, the juvenile vegetative phase is associated with small, round leaves that lack both trichomes on the abaxial leaf surface and serrations, whereas the adult phase is characterized by larger, elongated and serrated leaves that produce abaxial trichomes.

The core genetic network that controls the timing of vegetative phase change has been well described. The microRNA miR156, and its sister miR157, function as master regulators of the juvenile phase. A temporal decline in miR156/miR157 during shoot development leads to an increase in expression of their target genes—*SQUAMOSA PROMOTER BINDING PROTEIN-LIKE* (*SPL*) transcription factors—which promote the adult phase (Wu and Poethig, 2006; Wu et al., 2009). This temporal mechanism is widely conserved and regulates shoot identity in diverse plant lineages (Chuck et al., 2007; Leichty and Poethig, 2019; Riese et al., 2008; Wang et al., 2011). *SPL* genes are known to promote the expression of miR172, which initiates adult development through repression of its targets in the *APETALA2-LIKE* (*AP2-LIKE*) gene family. Vegetative phase change is thus promoted by inverse gradients of expression of two miRNAs, miR156 and miR172 (Wu et al., 2009).

*ALTERED MERISTEM PROGRAM1* (*AMP1*), which encodes a putative carboxypeptidase (Helliwell et al., 2001), was identified in a genetic screen for phase change mutations over 20 years ago (Conway and Poethig, 1997), but the basis for its effect on this process is still unknown. Mutations in *AMP1* produce a large number of small, round leaves that lack abaxial trichomes (juvenile leaves) and have a higher rate of leaf initiation (Telfer et al., 1997). An initial study suggested that this phenotype was not associated with a change in the timing of vegetative phase change, leading to the conclusion that the timing of vegetative phase change is regulated independently of leaf number (Telfer et al., 1997). However, this result conflicts with more recent studies showing that pre-existing leaves promote vegetative phase change (Yang et al., 2011; Yang et al., 2013; Yu et al., 2013). The phenotype of *amp1* is also surprising given the evidence that AMP1 is required for miRNA-mediated translational repression (Li et al., 2013). miR156 promotes juvenile development by translationally repressing its targets (He et al., 2018). If AMP1 is required for miRNA-mediated translational repression, *amp1* mutants would therefore be expected to have to a reduced number of juvenile leaves due to elevated *SPL* gene expression, which is the exact opposite of the *amp1* phenotype.

To resolve these issues, we investigated the interaction between *AMP1* and the miR156-*SPL* module. Our results indicate that *AMP1* promotes adult leaf traits in parallel to, or downstream of, the miR156-*SPL* module. We also found no evidence that AMP1 is required for translational repression by either miR156 or miR159. This latter result suggests that the mechanism by which miRNAs repress translation in plants is different for different transcripts.

## RESULTS AND DISCUSSION

### Elevated SPL activity has a modest effect on the *amp1* phenotype

*amp1-1* (hereafter, *amp1*) mutants resemble plants with reduced *SPL* gene expression in having an increased rate of leaf initiation, an increased number of rosette leaves, an enlarged shoot apical meristem, and small, round rosette leaves that lack abaxial trichomes (Fig. 1A-F) (Chaudhury et al., 1993; Huang et al., 2015; Telfer et al., 1997; Yang et al., 2018). To determine if this phenotype is attributable to a reduction in *SPL* activity, we introduced *35S::MIM156* — which de-represses *SPL* gene expression (Franco-Zorrilla et al., 2007) — into *amp1*. *35S::MIM156* plants have a relatively slow rate of leaf initiation, have enlarged and somewhat elongated rosette leaves, produce abaxial trichomes unusually early in shoot development, and have a relatively small SAM (Fig. 1A-F). *amp1; 35S::MIM156* plants had a vegetative phenotype intermediate between that of the two parental genotypes, but which was more similar to *amp1* than to *35S::MIM156*. The rosette leaves of *amp1; 35S::MIM156* were approximately the same size as *amp1* leaves, but were similar in shape to *35S::MIM156* (Fig. 1A, B). *amp1* plants rarely produced rosette leaves with abaxial trichomes (although abaxial trichome production on cauline leaves was unaffected (Fig. S1)), whereas about 25% of *amp1; 35S::MIM156* produced rosette leaves with abaxial trichomes late in shoot development. In contrast, all *35S::MIM156* plants produced rosette leaves with abaxial trichomes by plastochron 3 (Fig. 1C). Similarly, the rate of leaf initiation in *amp1; 35S::MIM156* was intermediate between that of *amp1* and *35S::MIM156*, but was closer to that of *amp1* than *35S::MIM156* (Fig. 1D). The number of rosette leaves in *amp1; 35S::MIM156* was also intermediate between these two genotypes, but was more similar to *amp1* than *35S::MIM156* (Fig. 1E). Finally, the SAM of *amp1; 35S::MIM156* was more similar in size to *amp1* than to *35S::MIM156* (Fig. 1F). These results suggest that the phenotype of *amp1* is not a consequence of repressed SPL activity, implying that *AMP1* acts either downstream or in parallel to the miR156/*SPL* module. This conclusion is consistent with the observation that vegetative development (Fig. 1C) and flowering time (Fig. 1G) are dissociated in *amp1*.

**Fig. 1.**
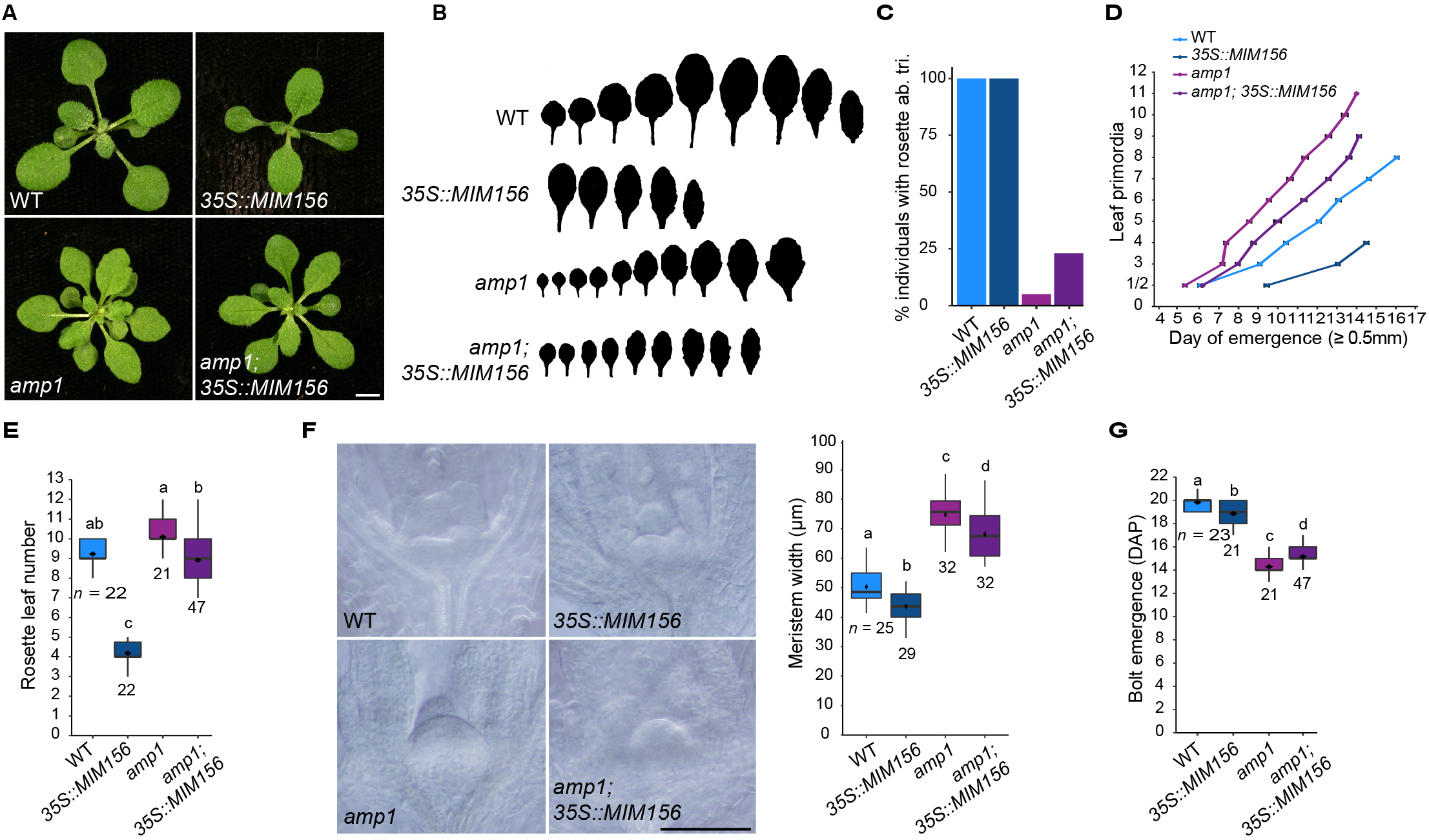
Elevated *SPL* gene activity only partially suppresses the *amp1* phenotype. (A) Photographs of plants taken at 16 DAP, scale bar: 5mm. (B) Silhouettes of heteroblastic series of rosette leaves for lines shown in (A). (C) Percentage of individual plants that produced at least one rosette leaf with abaxial trichomes (*n* ≥ 18). (D) Leaf emergence was scored when leaves became visible without manipulation of the rosette, error bars represent the SEM (*n* ≥ 18). (E, F, G) Statistically distinct genotypes were identified by one-way ANOVA with *post hoc* Tukey multiple comparison test (letters indicate statistically distinct groups; *p* < 0.05; sample sizes are shown in the figure). Images and measurements in (F) are of SAMs of plants harvested at 5 DAP captured using DIC microscopy, scale bar: 100μM. All phenotypic analyses were carried out in LD conditions.

### The phenotype of *amp1* is not attributable to a change in miR156/miR157 or *SPL* gene expression

To explore the relationship between *AMP1* and the miR156/*SPL* module in more detail, we examined the effect of *amp1* on the abundance of the miR156 and *SPL* transcripts. qRT-PCR analysis of the shoot apices of plants grown in short days (SD) showed that *amp1* had no significant effect on the level of miR156 or miR157 (Fig. 2A), or the transcripts of three direct targets of these miRNAs: *SPL3*, *SPL9* and *SPL13* (Fig. 2B). To test whether *amp1* affects *SPL* expression independent of miR156/miR157, we measured the transcripts of these genes in *35S::MIM156* and *amp1; 35S:MIM156* plants. As expected (He et al., 2018), all three *SPL* transcripts were significantly elevated in *35S::MIM156.* All three transcripts were elevated to a much smaller extent in *amp1; 35S:MIM156* (Fig. 2B). Together, these results suggest that *AMP1* may promote *SPL* expression, but only in the absence of miR156/miR157.

**Fig. 2.**
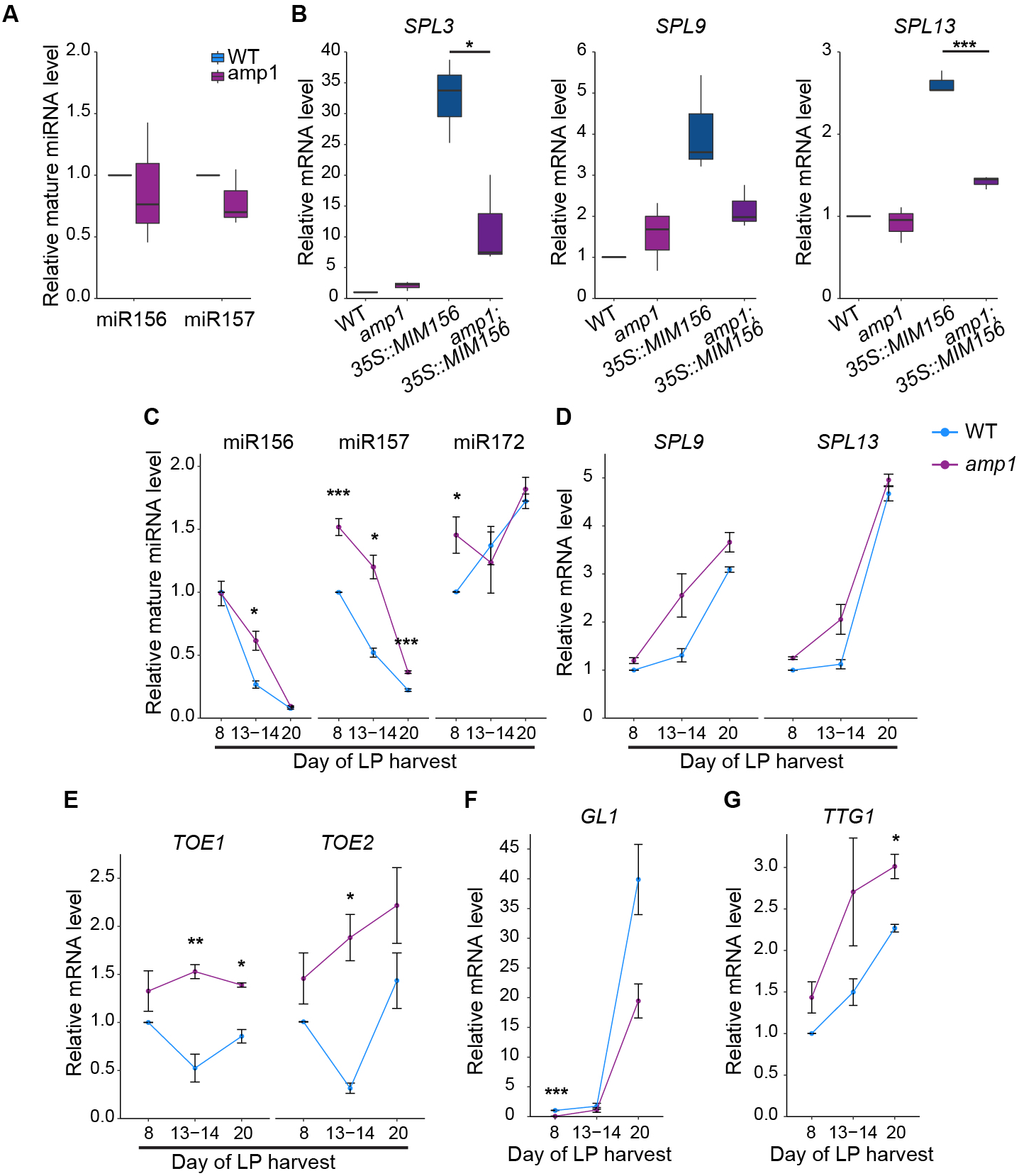
The *amp1* phenotype is not associated with repressed *SPL* activity. qRT-PCR analyses of gene expression. (A, B) Shoot apices with leaf primordia ≥ 1mm removed at 8 DAP. (C - G) Isolated leaf primordia (LP) 0.5-1mm in size. 8 DAP = LP1-2; 13-14 DAP = LP 4-5 (*amp1* LP were harvested at 13 DAP, WT LP at 14 DAP); 20 DAP = WT LP9-10, *amp1* LP14-16. Error bars represent the SEM. All plants were grown in SD conditions. Asterisks represent significant differences between genotypes calculated by two-tailed *t*-test (*p* < 0.05 = *; *p* < 0.01 = **; *p* < 0.001 = ***).

We then examined the expression of these genes in successive rosette leaf primordia (LP) of plants grown in SD. Because *amp1* initiates leaves more rapidly than WT, LP were grouped according to the time of harvest rather than position on the shoot. Both the level and rate of decline of miR156 were almost identical in WT and *amp1* (Fig. 2C). miR157 was elevated in all LP, but declined at approximately the same rate as in WT plants. *SPL9* and *SPL13* transcripts were also elevated in the LP of *amp1* relative to WT (Fig. 2D), but these differences were relatively modest (two-fold or less) and not statistically significant. Furthermore, the elevated expression of *SPL9* and *SPL13* is inconsistent with the elevated level of miR157 and with the juvenilized phenotype of *amp1*. Taken together, these data suggest that the vegetative phenotype of *amp1* is not caused by increased expression of miR156/miR157 or decreased expression of *SPL* genes. It is possible that *AMP1* regulates *SPL* expression independently of miR156 (Fig. 2B). However, the observation that *amp1* does not have a significant effect on *SPL9* and *SPL13* expression at 20 DAP (Fig. 2D), when the levels of miR156 and miR157 are very low (Fig. 2C), suggests that this is unlikely.

If *AMP1* does not regulate miR156 or *SPL* gene expression, perhaps it regulates shared downstream targets. Consistent with this hypothesis, expression of the closely-related *AP2-like* transcription factors *TOE1* and *TOE2* (which are targets of the SPL-regulated miRNA, miR172) was consistently elevated in *amp1* (Fig. 2E). This effect is not attributable to a change in the level of miR172, however, as the abundance of this miRNA was not reduced in *amp1* (Fig. 2C). *TOE1* blocks the production of trichomes on the abaxial side of the leaf by working in association with the abaxial specification gene *KANAD1* (*KAN1*) to repress the transcription of *GLABRA1* (*GL1*) (Wang et al., 2019; Xu et al., 2019). In WT plants, *GL1* expression increased dramatically between 13-14 DAP and 20 DAP, consistent with the increase in trichome production over this period. *GL1* displayed a similar temporal pattern in *amp1*, but was almost completely suppressed in the earliest LP and was considerably lower than WT in LP harvested at DAP (Fig. 2F). In contrast, the expression of *TRANSPARENT TESTA GLABRA1* (*TTG1*) — which promotes trichome initiation via a distinct protein complex to *GL1* (Pesch et al., 2015) — was not reduced in *amp1* (Fig. 2G). These results suggest that *AMP1* promotes abaxial trichome formation via *GL1*, not *TTG1*, and that it acts as a general activator of *GL1* expression, rather than a temporal regulator. They also support the conclusion that *AMP1* regulates abaxial trichome production downstream of miR156/*SPL*.

### The timing of vegetative phase change is regulated independently of leaf initiation in *amp1*

The juvenilized phenotype of *amp1* was originally attributed to the increased rate of leaf initiation in this mutant (Telfer et al., 1997). However, this interpretation is inconsistent with more recent studies showing that pre-existing leaves promote the transition to the adult vegetative phase by repressing miR156 (Yang et al., 2011; Yang et al., 2013; Yu et al., 2013). To determine the basis of this discrepancy, we characterized the effect of *CLAVATA3* (*CLV3*) and *CLV1* mutations on vegetative phase change. We chose these mutations because they resemble *amp1* in having an enlarged SAM and an accelerated rate of leaf initiation (Clark et al., 1995; Leyser and Furner, 1992).

Like *amp1* (Telfer et al., 1997), *clv3* and *clv1* produced smaller, rounder rosette leaves, and more leaves without abaxial trichomes (Fig. 3A - C). This increase in the number of juvenile-like leaves was not associated with a delay in the juvenile-to-adult transition, however. Instead, *clv3* mutants produced leaves with abaxial trichomes one day earlier than WT plants (Fig. 3D). To determine if the phenotype of *clv1* and *clv3* is dependent on miR156, we introduced the miR156 sponge, *35S::MIM156*, into these mutants. This transgene was epistatic to *clv1* and *clv3* with respect to their effect on leaf shape (Fig. 3A, B) and abaxial trichome production (Fig. 3C), suggesting that their effect on these traits requires miR156.

**Fig. 3.**
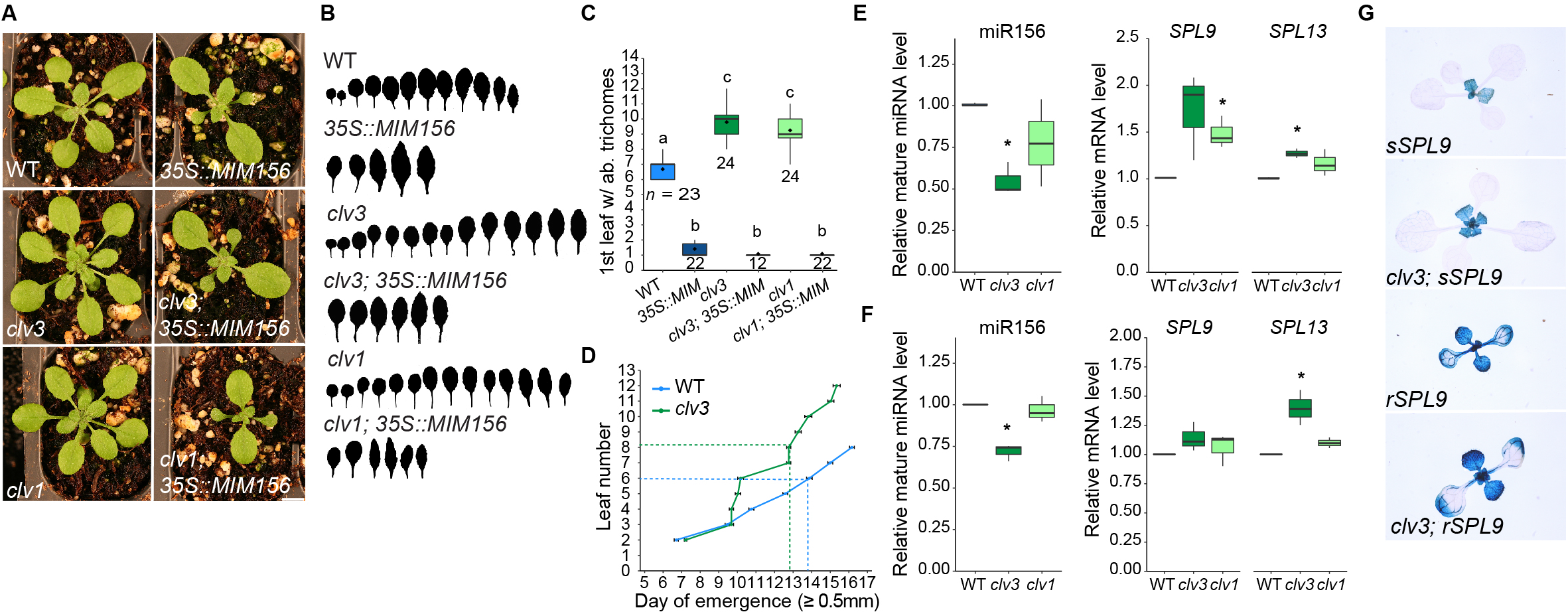
Enhanced *clv* juvenility is a consequence of an increased rate of leaf initiation, rather than repressed *SPL* gene activity. (A) Photographs were taken at DAP, scale bar: 5mm. (B) Silhouettes of heteroblastic series of rosette leaves for lines shown in (A). (C) Statistically distinct genotypes were identified by one-way ANOVA with *post hoc* Tukey multiple comparison test (letters indicate statistically distinct groups; *p* < 0.05; sample sizes are shown in the figure). (D) Leaf emergence was scored when leaves became visible without manipulation of the rosette, the dashed line indicates the first leaf to produce abaxial trichomes, error bars represent the SEM (*n* ≥ 23). (E, F) qRT-PCR analyses of gene expression of (E) shoot apices with leaf primordia ≥ 1mm removed at 10 DAP and (F) isolated LP1&2 0.5-1mm in size. Asterisks represent significant differences between WT vs *clv3* or WT vs *clv1*. Significance was calculated by two-tailed *t*-test (*p* < 0.05). (G) GUS staining of miR156-sensitive and miR156-resistant SPL-GUS reporter constructs at 14 DAP. Phenotypic analyses were carried out in LD conditions (A-D), gene expression analyses were carried out in SD conditions (E-G).

We then examined the effect of *clv3* and *clv1* on the expression of miR156 and its targets, *SPL9* and *SPL13*, in shoot apices (Fig. 3E) and LP (Fig. 3F). qRT-PCR revealed that *clv1* and *clv3* have slightly reduced levels of miR156, although this difference was only statistically significant in *clv3*. Consistent with the decreased amount of miR156, *SPL9* and *SPL13* transcripts were slightly elevated in both the mutants, although again this difference was only statistically significant in a few cases. If these relatively small differences in miR156 and *SPL* gene expression are functionally significant, they would be expected to promote the appearance of adult traits, not repress the expression of these traits as is the case in *clv1* and *clv3*. To explore this inconsistency, we examined the effect of *clv3* on the expression of a miR156-sensitive and a miR156-resistant version of the *SPL9::SPL9-GUS* reporter (Xu et al., 2016). There was no obvious difference in the expression of these reporters in the presence or absence of *clv3* (Fig. 3G), supporting the conclusion that the effect of *clv3* on leaf identity is not attributable to a change in the level of miR156 or its targets.

Instead, the effect of *clv3* and *clv1* on leaf identity is primarily attributable to their effect on the rate of leaf initiation. Specifically, *clv3* and *clv1* appear to increase the number of juvenile leaves by accelerating the rate of leaf production during the period when miR156 levels are high. This conclusion is supported by the observation that *35S::MIM156* is epistatic to these mutations with respect to their effect on leaf identity (Fig. 3A, B); i.e. miR156 is required for their leaf identity phenotypes. Consistent with the evidence that leaves promote the juvenile-to-adult transition by repressing miR156 (Yang et al., 2011; Yang et al., 2013; Yu et al., 2013), *clv3* and *clv1* have slightly reduced levels of miR156 and slightly elevated levels of *SPL9* and *SPL13* (Fig. 3E, F). However, this relatively small effect is apparently insufficient to interfere with the function of these genes in specifying juvenile leaf identity.

The increased number of juvenile leaves in *amp1* is also partly attributable to its higher rate of leaf initiation (Telfer et al., 1997). However, *amp1* differs from *clv3* and *clv1* in having a much more significant effect on leaf identity: *amp1* rarely produces abaxial trichomes on rosette leaves, whereas *clv3* and *clv1* routinely do so. In addition, the phenotype of *amp1* is less sensitive to a reduction in miR156 than the phenotype of *clv3* and *clv1*; in general, *amp1, 35S::MIM156* plants more closely resembled *amp1* than *35S::MIM156* (Fig. 1A-E). This observation, and the effect of *amp1* on the expression of genes involved in abaxial trichome production (Fig. 2E, F), suggest that AMP1 operates independently of miR156 to regulate genes involved in leaf identity. A direct effect of AMP1 on leaf identity genes would explain why *amp1* has a more severe vegetative phenotype than *clv3* and *clv1*, and why the phenotype of *amp1* is relatively insensitive to changes in the level of miR156.

### *AMP1* is not universally required for translational repression

Given the role of *AMP1* in translational repression (Li et al., 2013), it is possible that the abundance of *SPL* transcripts in *amp1* (Fig. 2B, D) does not accurately reflect their biological activity. To determine whether AMP1 is required for the post-transcriptional regulation of *SPL* genes, we first measured the amount of *SPL9* and *SPL13* transcript cleavage in WT and *amp1* plants. Consistent with a previous study on miR156-mediated cleavage (He et al., 2018), the rate of transcript cleavage for both *SPL9* and *SPL13* declined during vegetative development in WT plants (Fig. 4A). This happened at a slower rate in *amp1*, presumably in part due to the higher level of miR156 in the *amp1* 13-14 DAP sample compared to WT (Fig. 2C) and the threshold-dependence of miR156 activity (He et al., 2018). However, later in development, transcript cleavage in *amp1* was similar to WT (Fig. 4A). This demonstrates that miR156 is functional in *amp1* and confirms the observation that AMP1 is not required for transcriptional cleavage (Li et al., 2013).

**Fig. 4.**
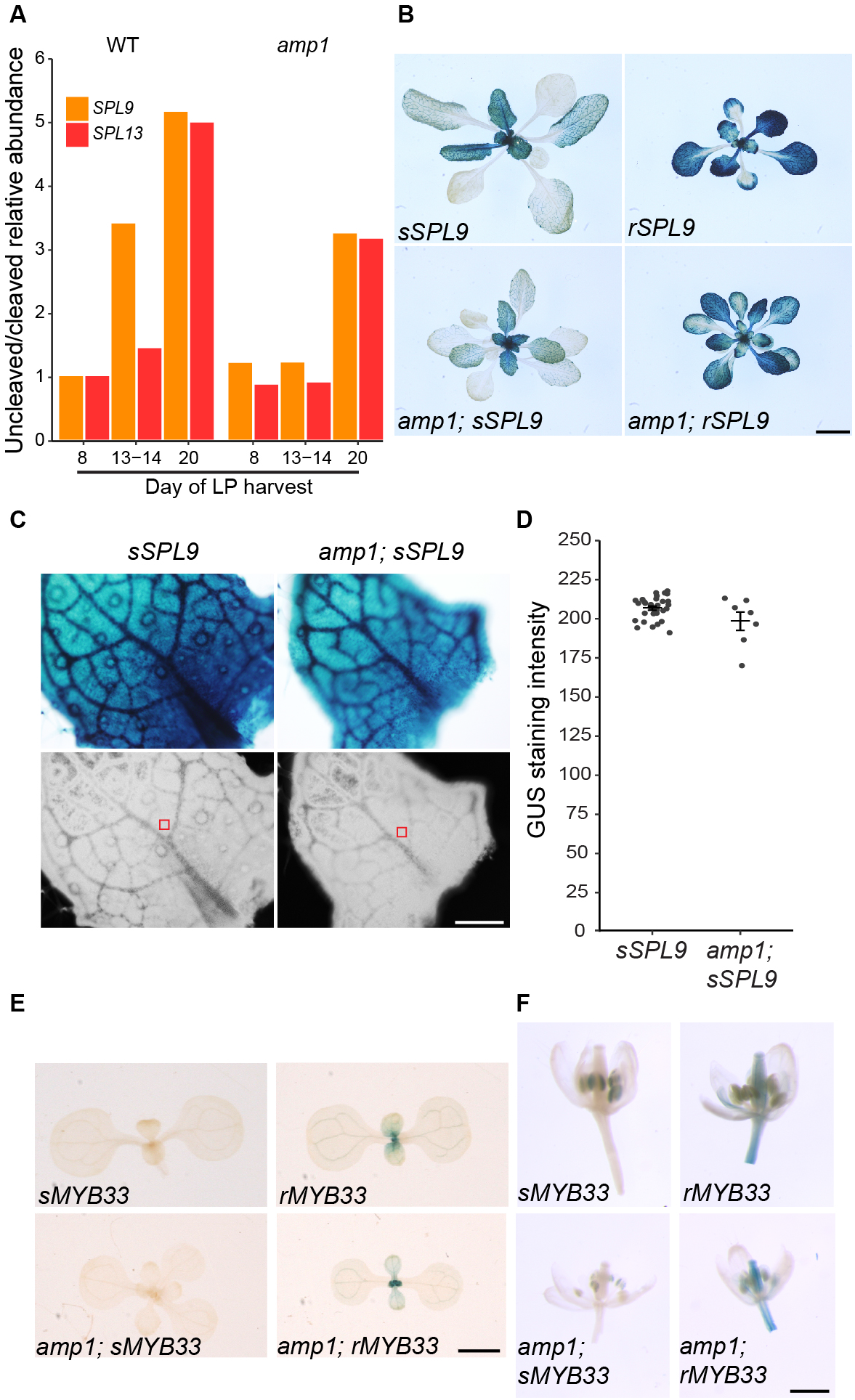
miRNA-regulated SPL9 and MYB33 proteins accumulate normally in *amp1*. (A) The relative abundance of uncleaved/cleaved transcripts, normalized to WT 8 DAP. See Fig. 2B legend for details of samples. (B) GUS staining of miR156-sensitive and miR156-resistant SPL-GUS reporter constructs at 21 DAP. (C, D) Quantification of sSPL9-GUS protein levels by image analysis. RGB color mode ((C), top panels), hue saturation brightness mode ((C), bottom panels). Red squares indicate where signal intensity was measured, each dot represents an individual primordia (D). (E, F) GUS staining of miR159-sensitive and miR159-resistant MYB33-GUS reporter constructs in 7 DAP seedlings (E) and flowers (F). Scale bars: (B) 5mm, (C) 200μM, (E, F) 1mm.

Although miR156 induces transcript cleavage, it represses the expression of its targets primarily by promoting translational repression (He et al., 2018). To examine the effect of *amp1* on this process, we crossed miR156-sensitive (sSPL9) and miR156-resistant (rSPL9) GUS-reporter constructs of SPL9 into *amp1*. There was no obvious difference in the staining intensity of these reporter proteins in WT and *amp1* (Fig. 4B). To confirm this impression, we measured the staining intensity of the sSPL9-GUS reporter spectrophotometrically in leaf primordia of WT and *amp1* harvested at a stage when transcript cleavage was nearly equivalent in these genotypes (20 DAP (Fig. 4A)). There was no significant difference in sSPL9 protein levels in these genotypes (Fig. 4C, D). These results indicate that *amp1* has no effect on the activity of miR156, implying that translational repression of *SPL9* occurs normally in *amp1*. To determine if miR156 is uniquely insensitive to *amp1*, we examined the effect of *amp1* on the expression of MYB33, a transcription factor that also regulates shoot identity (Guo et al., 2017) and is translationally repressed by miR159 (Li et al., 2014). miR159-sensitive and miR159-resistant versions of MYB33-GUS (Millar and Gubler, 2005) were crossed into *amp1*, and WT and *amp1* plants were stained for GUS activity one week after germination, and at flowering. MYB33-GUS was repressed in a miR159-dependent fashion in leaves and floral organs of WT plants, and *amp1* had no obvious effect on this expression pattern (Fig. 4E, F). Because *amp1* had no effect on the expression of sMYB33-GUS, it is reasonable to assume that miR159-dependent translational repression occurs normally in this mutant. We conclude from these results that *AMP1* is not universally required for the translational repression of miRNA-targets.

Whether or not *AMP1* functions in translational repression may be a result of the sub-cellular localization of the process. AMP1 has been shown to colocalize with the key silencing component ARGONAUTE1 (AGO1) on the endoplasmic reticulum (ER) (Li et al., 2013). However, AGO1 also localizes to processing bodies (p-bodies), cytoplasmic foci of mRNA-ribonucleoprotein complexes that facilitate the sequestration of mRNAs for translational silencing (reviewed in Chantarachot and Bailey-Serres, 2018). Loss of the p-body protein SUO leads to a reduction in the translational repression of the miR156-target *SPL3* (Yang et al., 2012), suggesting that p-bodies are also important sites of miRNA-mediated translational repression. Taken together, these results are consistent with a model in which a) miRNA-mediated translational repression occurs in distinct sub-cellular compartments in a sequence-specific manner and b) unique sets of proteins contribute to this repression, depending on the compartment (e.g. AMP1 on the ER, SUO in p-bodies). Whether the translational repression of *MYB33* by miR159 occurs in p-bodies remains to be demonstrated.

Support for this model comes from the finding that the microtubule severing-enzyme KATANIN 1 is also required for translation repression (Brodersen et al., 2008). What signals the cellular machinery uses to determine where to localize miRNA-target pairs for translational repression is unclear. There appear to be no consistent differences between the miRNA hairpin secondary structures and miRNA/miRNA* duplexes of AMP1-dependent and AMP1-independent miRNAs (Fig. S2). Although it is perhaps unlikely that any such signals would persist during miRNA processing. The strength of target complementarity is known to affect silencing efficacy (Li et al., 2014), and could also drive sub-cellular distribution, but there is also no trend in target mismatch number between the AMP1-dependent/independent classes of miRNA (Table S1). Given the overlapping expression domains of a number of these miRNAs (reviewed in Fouracre and Poethig, 2016), it is unlikely that the site of translational repression is developmentally regulated. At the cellular level, there is evidence to suggest that miRNA sequences include signals that control the specificity of inter-cellular mobility (Skopelitis et al., 2018). It will be fascinating to see if the same signaling mechanisms determine the destination of miRNAs within cells.

## Materials and Methods

### Plant material and growth conditions

Col was used as the genetic background for all stocks. The following genetic lines have been described previously: *amp1-1* (Chaudhury et al., 1993); *SPL9::sSPL9-GUS, SPL9::rSPL9-GUS* (Xu et al., 2016); *35S::MIM156* (Fouracre and Poethig, 2019); *clv1-4* (Clark et al., 1993); *MYB33::sMYB33-GUS, MYB33::rMYB33-GUS* (Millar and Gubler, 2005). *clv3-10* (CS68823) was obtained from the Arabidopsis Biological Resource Center (Ohio State University). Seeds were sown on fertilized Farfard #2 soil (Farfard) and kept at 4°C for 3 days prior to transfer to a growth chamber, with the transfer day counted as day 0 for plant age (0 DAP). Plant were grown at 22°C under a mix of both white (USHIO F32T8/741) and red-enriched (Interlectric F32/T8/WS Gro-Lite) fluorescent bulbs in either long day (16 hrs. light/8 hrs. dark; 80 μmol m^−2^ s^−1^) or short day (10 hrs light/14 hrs dark; 120 μmol m^−2^ s^−1^) conditions.

### GUS staining

Plants were fixed in 90% acetone on ice for 10 minutes and washed with GUS staining buffer (5mM potassium ferricyanide and 5mM ferrocyanide in 0.1M PO_4_ buffer) and stained for between 8 hrs and overnight (depending on transgene strength) at 37°C in 2mM X-Gluc GUS staining buffer. For the quantification of GUS staining intensity, ~1mm LP were harvested at 21 DAP, stained O/N and images of stained primordia converted from RGB color mode to hue saturation brightness mode as previously described (Béziat et al., 2017). A consistent position in the middle of the leaf lamina, adjacent to the midvein, was used for measurement.

### Histology

Shoot apices were cleared and imaged according to a described protocol (Chou et al., 2016).

### RNA expression analyses

Tissue (either shoot apices with leaf primordia ≤1mm attached or isolated leaf primordia 0.5-1mm in size – as specified in the text) were ground in liquid nitrogen and total RNA extracted using Trizol (Invitrogen) as per the manufacturer’s instructions. RNA was DNAse treated with RQ1 (Promgea) and 250ng-1μg of RNA was used for reverse transcription using Superscript III (Invitrogen). Gene specific RT primers were used to amplify miR156, miR157, miR172 and SnoR101 and a polyT primer for mRNA amplification. Three-step qPCR of cDNA was carried out using SYBR-Green Master Mix (Bimake). qPCR reactions were run in triplicate and an average taken. For analyses of *amp1* shoot apices and *clv* mutants, separate RNA extractions of three biological replicates were carried out. For analyses of *amp1* leaf primordia, three reverse-transcription replicates from single RNA extractions were carried out for each sample (at least 60 LP were pooled for each RNA extraction). 8 DAP samples were collected twice - once as part of a biological replicate with 13-13 DAP and once as part of a biological replicate with 20 DAP samples. Relative transcript levels were normalized to snoR101 (for miRNAs) and *ACT2* (*amp1* shoot apices, *clv* mutants) or *UBQ10* (*amp1* leaf primordia) (for mRNAs) and expressed as a ratio of expression to WT (*amp1* shoot apices, *clv* mutants) and WT 8 DAP (*amp1* leaf primordia) samples

For the quantification of transcript cleavage, a modified 5’RACE protocol was followed as previously described (He et al., 2018). The data presented are the average of three ratios from separate reverse transcription replicates (six in the case of *amp1* 8 DAP – three reverse transcription replicates from two biological replicates).

The qPCR primers used in this study are listed in Supplementary Table 2.

### Statistical analyses

A two-tailed Student’s *t*-test was used to carry out pairwise comparisons between different genotypes. For comparison of multiple samples, to decrease the chance of false positives, a one-way ANOVA followed by a Tukey test was used for multi-way comparisons. Statistical analyses were carried out in R (r-project.org) and Excel (Microsoft).

For figures featuring boxplots, boxes display the IQR (boxes), median (lines), and values beyond 1.5* IQR (whiskers); mean values are marked by a solid diamond (♦).

## Acknowledgements

We thank Anthony Millar (Australian National University) for the kind gift of the MYB33- GUS reporters, members of the Poethig lab for useful discussions and Melissa Morrison for assistance with collecting phenotypic data.

## Figure Legends

**Supplementary Fig. 1. *amp1* cauline leaves produce abaxial trichomes.** Scale bar: 2mm

**Supplementary Fig. 2 Predicted hairpin structures for miRNAs that are *AMP1*-dependent and independent for translational repression.** *AMP1*-independent (this study) – miR156, miR157, miR159; *AMP1*-dependent (Li et al., 2013) – miR164, miR165, miR166 and miR398. Representative functional members of miRNA families are displayed. Stem-loop sequences were downloaded from miRBase (www.mirbase.org) and hairpin structures predicted using the default settings on RNAfold (http://rna.tbi.univie.ac.at/). Minimum free-energy models of hairpins are shown, color coded for base pair probability. Black lines are drawn alongside mature miRNA sequences.

## References

Béziat, C., Kleine-Vehn, J. and Feraru, E. (2017). Histochemical Staining of β-Glucuronidase and Its Spatial Quantification. In Plant Hormones: Methods and Protocols (ed. Kleine-Vehn, J. and Sauer, M.), pp. 73–80. New York, NY: Springer New York.

Brodersen, P., Sakvarelidze-Achard, L., Bruun-Rasmussen, M., Dunoyer, P., Yamamoto, Y. Y., Sieburth, L. and Voinnet, O. (2008). Widespread translational inhibition by plant miRNAs and siRNAs. Science 320, 1185–1190.

Chantarachot, T. and Bailey-Serres, J. (2018). Polysomes, Stress Granules, and Processing Bodies: A Dynamic Triumvirate Controlling Cytoplasmic mRNA Fate and Function. Plant Physiol. 176, 254–269.

Chaudhury, A. M., Letham, S., Craig, S. and Dennis, E. S. (1993). *amp1* - a mutant with high cytokinin levels and altered embryonic pattern, faster vegetative growth, constitutive photomorphogenesis and precocious flowering. The Plant Journal 4, 907–916.

Chou, H., Wang, H. and Berkowitz, G. A. (2016). Shoot apical meristem size measurement. Bio-protocol 6, e2055.

Chuck, G., Cigan, A. M., Saeteurn, K. and Hake, S. (2007). The heterochronic maize mutant *Corngrass1* results from overexpression of a tandem microRNA. Nat Genet 39, 544–549.

Clark, S. E., Running, M. P. and Meyerowitz, E. M. (1993). *CLAVATA1*, a regulator of meristem and flower development in *Arabidopsis*. Development 119, 397–418.

Clark, S. E., Running, M. P. and Meyerowitz, E. M. (1995). *CLAVATA3* is a specific regulator of shoot and floral meristem development affecting the same processes as *CLAVATA1* Development 121, 2057–2067.

Conway, L. J. and Poethig, R. S. (1997). Mutations of *Arabidopsis thaliana* that transform leaves into cotyledons. Proc Natl Acad Sci USA 94, 10209–10214.

Fouracre, J. P. and Poethig, R. S. (2016). The role of small RNAs in vegetative shoot development. Curr Opin Plant Biol 29, 64–72.

Fouracre, J. P. and Poethig, R. S. (2019). Role for the shoot apical meristem in the specification of juvenile leaf identity in *Arabidopsis*. Proc Natl Acad Sci USA 116, 10168.

Franco-Zorrilla, J. M., Valli, A., Todesco, M., Mateos, I., Puga, M. I., Rubio-Somoza, I., Leyva, A., Weigel, D., Garcia, J. A. and Paz-Ares, J. (2007). Target mimicry provides a new mechanism for regulation of microRNA activity. Nat Genet 39, 1033–7.

Guo, C., Xu, Y., Shi, M., Lai, Y., Wu, X., Wang, H., Zhu, Z., Poethig, R. S. and Wu, G. (2017). Repression of miR156 by miR159 regulates the timing of the juvenile-to-adult transition in *Arabidopsis*. Plant Cell 29, 1293–1304.

He, J., Xu, M., Willmann, M. R., McCormick, K., Hu, T., Yang, L., Starker, C. G., Voytas, D. F., Meyers, B. C. and Poethig, R. S. (2018). Threshold-dependent repression of *SPL* gene expression by miR156/miR157 controls vegetative phase change in *Arabidopsis thaliana*. PLoS Genet. 14, e1007337.

Helliwell, C. A., Chin-Atkins, A. N., Wilson, I. W., Chapple, R., Dennis, E. S. and Chaudhury, A. (2001). The *Arabidopsis AMP1* gene encodes a putative glutamate carboxypeptidase. Plant Cell 13, 2115–2125.

Huang, W., Pitorre, D., Poretska, O., Marizzi, C., Winter, N., Poppenberger, B. and Sieberer, T. (2015). *ALTERED MERISTEM PROGRAM1* Suppresses Ectopic Stem Cell Niche Formation in the Shoot Apical Meristem in a Largely Cytokinin-Independent Manner. Plant Physiol 167, 1471–1486.

Huijser, P. and Schmid, M. (2011). The control of developmental phase transitions in plants. Development 138, 4117–4129.

Leichty, A. R. and Poethig, R. S. (2019). Development and evolution of age-dependent defenses in ant-acacias. Proc Natl Acad Sci USA 116, 15596.

Leyser, H. M. O. and Furner, I. J. (1992). Characterization of three shoot apical meristem mutants of *Arabidopsis thaliana*. Development 116, 397–403.

Li, S., Liu, L., Zhuang, X., Yu, Y., Liu, X., Cui, X., Ji, L., Pan, Z., Cao, X., Mo, B., et al. (2013). MicroRNAs inhibit the translation of target mRNAs on the endoplasmic reticulum in *Arabidopsis*. Cell 153, 562–74.

Li, J., Reichel, M. and Millar, A. A. (2014). Determinants beyond both complementarity and cleavage govern microR159 efficacy in *Arabidopsis*. PLoS Genet 10, e1004232.

Millar, A. A. and Gubler, F. (2005). The Arabidopsis GAMYB-like genes, *MYB33* and *MYB65*, are microRNA-regulated genes that redundantly facilitate anther development. Plant Cell 17, 705–21.

Pesch, M., Schultheiß, I., Klopffleisch, K., Uhrig, J. F., Koegl, M., Clemen, C. S., Simon, R., Weidtkamp-Peters, S. and Hülskamp, M. (2015). TRANSPARENT TESTA GLABRA1 and GLABRA1 Compete for Binding to GLABRA3 in *Arabidopsis*. Plant Physiol. 168, 584–597.

Poethig, R. S. (2013). Vegetative phase change and shoot maturation in plants. Curr. Top. Dev. Biol. 105, 125–52.

Riese, M., Zobell, O., Saedler, H. and Huijser, P. (2008). SBP-domain transcription factors as possible effectors of cryptochrome-mediated blue light signalling in the moss *Physcomitrella patens*. Planta 227, 505–15.

Skopelitis, D. S., Hill, K., Klesen, S., Marco, C. F., von Born, P., Chitwood, D. H. and Timmermans, M. C. P. (2018). Gating of miRNA movement at defined cell-cell interfaces governs their impact as positional signals. Nat. Commun. 9, 3107.

Telfer, A., Bollman, K. M. and Poethig, R. S. (1997). Phase change and the regulation of trichome distribution in *Arabidopsis thaliana*. Development 124, 645–654.

Wang, J. W., Park, M. Y., Wang, L. J., Koo, Y., Chen, X. Y., Weigel, D. and Poethig, R. S. (2011). miRNA control of vegetative phase change in trees. PLoS Genet 7, e1002012.

Wang, L., Zhou, C.-M., Mai, Y.-X., Li, L.-Z., Gao, J., Shang, G.-D., Lian, H., Han, L., Zhang, T.-Q., Tang, H.-B., et al. (2019). A spatiotemporally regulated transcriptional complex underlies heteroblastic development of leaf hairs in Arabidopsis thaliana. The EMBO Journal 38, e100063.

Wu, G. and Poethig, R. S. (2006). Temporal regulation of shoot development in *Arabidopsis thaliana* by miR156 and its target *SPL3*. Development 133, 3539–47.

Wu, G., Park, M. Y., Conway, S. R., Wang, J. W., Weigel, D. and Poethig, R. S. (2009). The sequential action of miR156 and miR172 regulates developmental timing in *Arabidopsis*. Cell 138, 750–9.

Xu, M., Hu, T., Zhao, J., Park, M. Y., Earley, K. W., Wu, G., Yang, L. and Poethig, R. S. (2016). Developmental functions of miR156-regulated *SQUAMOSA PROMOTER BINDING PROTEIN-LIKE* (*SPL*) genes in *Arabidopsis thaliana*. PLoS Genet. 12, e1006263.

Xu, Y., Qian, Z., Zhou, B. and Wu, G. (2019). Age-dependent heteroblastic development of leaf hairs in Arabidopsis. New Phytologist 224, 741–748.

Yang, L., Conway, S. R. and Poethig, R. S. (2011). Vegetative phase change is mediated by a leaf-derived signal that represses the transcription of miR156. Development 138, 245–9.

Yang, L., Wu, G. and Poethig, R. S. (2012). Mutations in the GW-repeat protein SUO reveal a developmental function for microRNA-mediated translational repression in Arabidopsis. Proc Natl Acad Sci USA 109, 315–320.

Yang, L., Xu, M., Koo, Y., He, J. and Poethig, R. S. (2013). Sugar promotes vegetative phase change in *Arabidopsis thaliana* by repressing the expression of *MIR156A* and *MIR156C*. Elife 2, e00260.

Yang, S., Poretska, O. and Sieberer, T. (2018). ALTERED MERISTEM PROGRAM1 Restricts Shoot Meristem Proliferation and Regeneration by Limiting HD-ZIP III-Mediated Expression of RAP2.6L. Plant Physiol. 177, 1580–1594.

Yu, S., Cao, L., Zhou, C. M., Zhang, T. Q., Lian, H., Sun, Y., Wu, J. Q., Huang, J. R., Wang, G. D. and Wang, J. W. (2013). Sugar is an endogenous cue for juvenile-to-adult phase transition in plants. Elife 2, e00269.

