## Supplemental Figures 1 and 2, Supplemental Tables 1 and 2 for "*ALTERED MERISTEM PROGRAM1* regulates leaf identity independent of miR156-mediated translational repression"

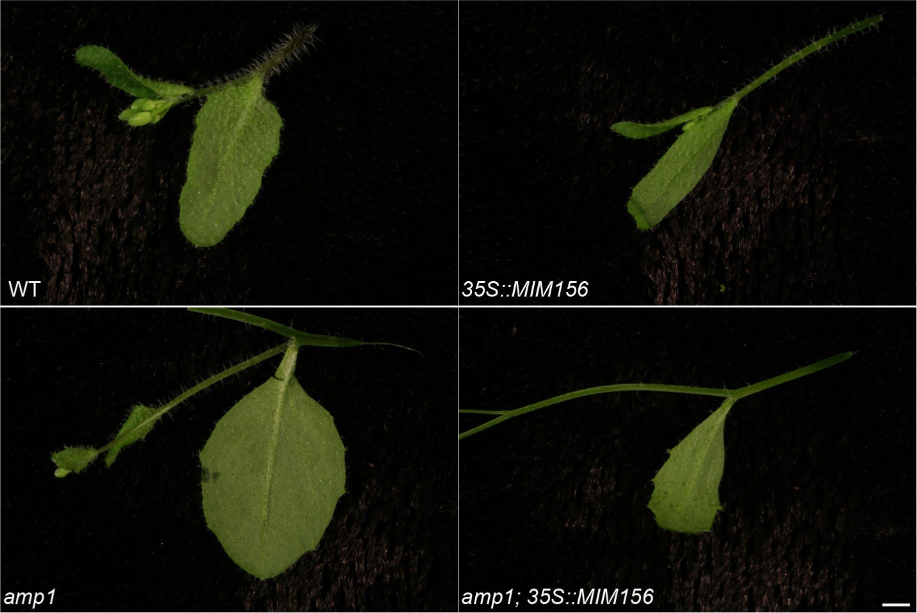
WT
35S::MIM156
amp1
amp1; 35S::MIM156

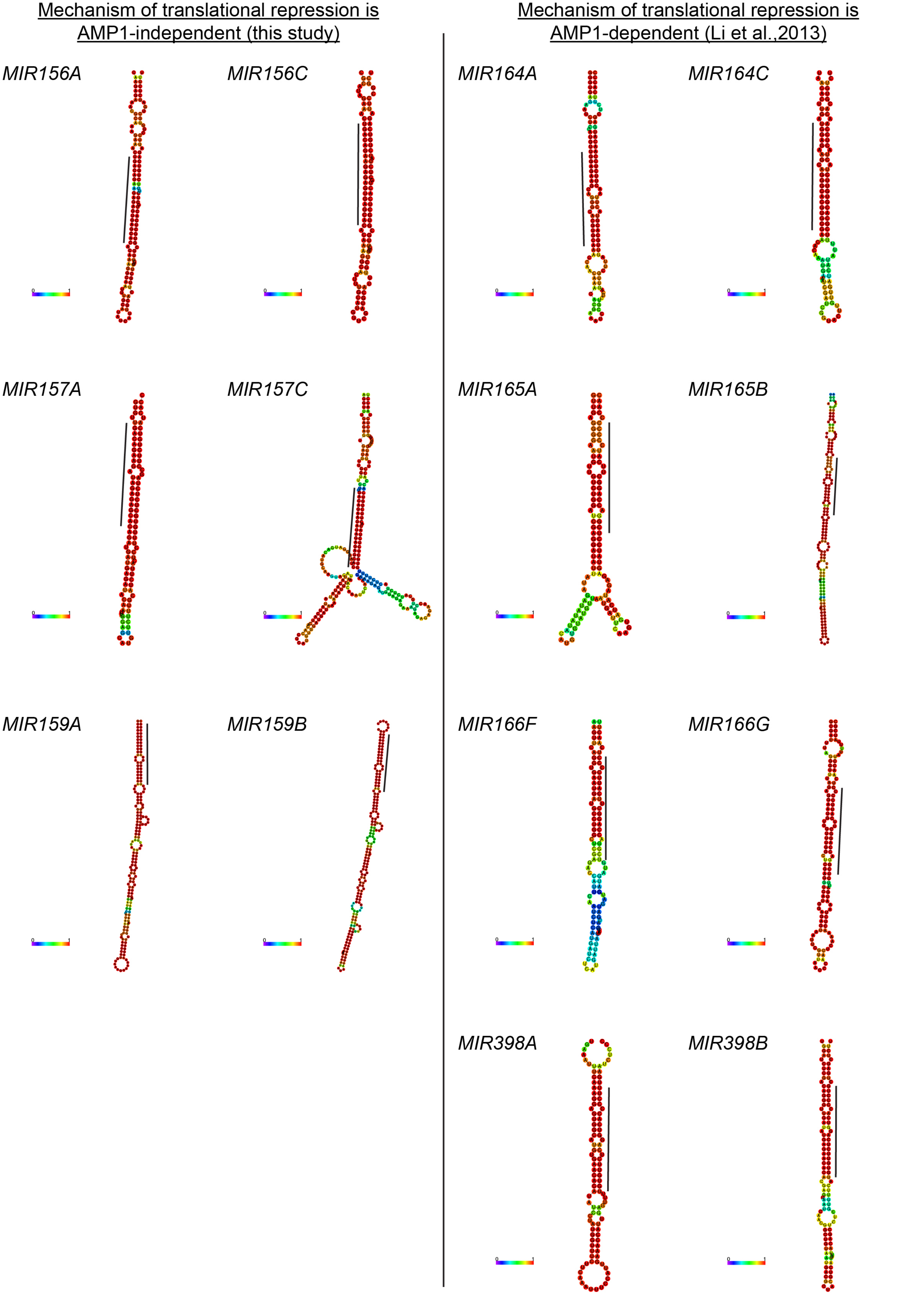
Mechanism of translational repression is AMP1-independent (this study)
Mechanism of translational repression is AMP1-dependent (Li et al.,2013)
MIR156A
MIR156C
MIR164A
MIR164C
MIR157A
MIR157C
MIR165A
MIR165B
MIR159A
MIR159B
MIR166F
MIR166G
MIR398A
MIR398B

| MicroRNA | Translational repression AMP1-dependent | Length (nt) | Duplex mismatch [insertion on star strand] [1] | Target mismatch [2] |
|---|---|---|---|---|
| *MIR156A* | No (this study) | 20 | [2] | 1 (*SPL9*) |
| *MIR156C* | No (this study) | 20 | [2] | 1 (*SPL9*) |
| *MIR157A* | No (this study) | 21 | 2 [1] | 1 (*SPL9*) |
| *MIR157C* | No (this study) | 21 | 1 [1] | 1 (*SPL9*) |
| *MIR159A* | No (this study) | 21 | 2 | 3 (*MYB33*) |
| *MIR159B* | No (this study) | 21 | 3 | 4 (*MYB33*) |
| *MIR164A* | Yes (Li et al 2013) | 21 | 3 | 3 (*CUC1*) |
| *MIR164C* | Yes (Li et al 2013) | 21 | 2 | 3 (*CUC1*) |
| *MIR165A* | Yes (Li et al 2013) | 21 | 4 | 3 (*PHB, REV*) |
| *MIR165B* | Yes (Li et al 2013) | 21 | 4 | 3 (*PHB, REV*) |
| *MIR166F* | Yes (Li et al 2013) | 21 | 4 | 4 (*PHB, REV*) |
| *MIR166G* | Yes (Li et al 2013) | 21 | 5 | 4 (*PHB, REV*) |
| *MIR398A* | Yes (Li et al 2013) | 21 | 3 | 6 (*CSD2*) |
| *MIR398B* | Yes (Li et al 2013) | 21 | 2 | 6 (*CSD2*) |

[1] Base pair mismatches not bracketed, un-paired insertions on star strand in brackets
[2] G::U wobbles included as mismatch

| | Primer name | Sequence |
|---|---|---|
| Reverse transcription | miR156 RT | GTCGTATCCAGTGCAGGGTCCGAGGTATTCGCACTGGATACGACGTGCTCA |
| | miR157 RT | GTCGTATCCAGTGCAGGGTCCGAGGTATTCGCACTGGATACGACGTGCTCT |
| | miR172 RT | GTCGTATCCAGTGCAGGGTCCGAGGTATTCGCACTGGATACGACATGCAG |
| | AtSnoR101 R1 | AGCATCAGCAGACCAGTAGTT |
| | Oligo dT | TTTTTTTTTTTTTTTTTTTTT |
| qPCR | ACT2-F | GCACCCTGTTCTTCTTACCG |
| | ACT2-R | AACCCTCGTAGATTGGCACA |
| | UBQ10-R | AAAGAGATAACAGGAACGGAAACATA |
| | UBQ10-F | GGCCTTGTATAATCCCTGATGAATAA |
| | qSPL9-F4 | AATTGGCGACTCAAACTGTG |
| | qSPL9-R4 | CTGAAGAAGCTCGCCATGTA |
| | qSPL13-F2 | GCTCGAGAACCGCATCGTT |
| | qSPL13-R2 | CCCGTAAAAAACTGTCTCAACTGCT |
| | qSPL-Cleaved-F1 | GGACTGAAGGAGTAGAAATCTTCTG |
| | qPCR SPL3 F | ATGAGTATGAGAAGAAGCAAAGCG |
| | qPCR SPL3 R | TCCACTACTACTTGTAGCTTTACCT |
| | QRT-TOE1-F | ACCGGTAATGCGCCAAAGCAAA |
| | QRT-TOE1-R | ACCGGCTTTCCCATGTATTCGT |
| | QRT-TOE2-F | TGCCCTTCCTTCTGCGTTCTTT |
| | QRT-TOE2-R | ACTGATCATGCCCTTGCCATGT |
| | GL1_F | GTGAACAAAGGCAATTTCACTG |
| | GL1_R | GTTCTTCCCGGTACTCTTTTAGC |
| | TTG1-qRTPCRf | GCGATTTCCTCCGTCTTTGG |
| | TTG1-qRTPCRr | CGCTCGTTTTGCTGTTGTTG |
| | AtSnoR101 F1 | CTTCACAGGTAAGTTCGCTTG |
| | AtSnoR101 R1 | AGCATCAGCAGACCAGTAGTT |
| | miR156 F | GCGGCGGTGACAGAAGAGAGT |
| | miR157 F | GCGGCGGTTGACAGAAGATAG |
| | miR172 F | CGGCGGAGAATCTTGATGATGC |
| | miRNA R | GTGCAGGGTCCGAGGT |
